# Shortening injection matrix for serial crystallography

**DOI:** 10.1101/687640

**Authors:** Ki Hyun Nam

**Affiliations:** Division of Biotechnology, Korea University, Seoul, Republic of Korea

**Keywords:** serial crystallography, SMX, shortening, delivery medium, sample delivery

## Abstract

Serial crystallography (SX) allows crystal structures to be observed at room temperature through the steady delivery of crystals to the X-ray interaction point. Viscous delivery media are advantageous because they afford efficient sample delivery from an injector or syringe at a low flow rate. Hydrophobic delivery media, such as lipidic cubic phase (LCP) or grease, provide a very stable injection stream and are widely used. The development of new hydrophobic delivery materials can expand opportunities for future SX studies with various samples. Here, I introduce fat-based shortening as a delivery medium for SX experiments. This material is commercially available at low cost and is straightforward to handle because its phase (i.e., solid or liquid) can be controlled by temperature. Shortening was extruded from a syringe needle in a very stable injection stream even below 200 nl/min. X-ray exposed shortening produced several background scattering rings, which have similar or lower intensities than those of LCP and contribute negligibly to data processing. Serial millisecond crystallography was performed using two shortening delivery media, and the room temperature crystal structures of lysozyme and glucose isomerase were successfully determined at resolutions of 1.5–2.0 Å. Therefore, shortening can be used as a sample delivery medium in SX experiments.

## Introduction

Serial crystallography (SX) using an X-ray free electron laser (XFEL) or synchrotron X-ray sources allows crystal structure to be determined without radiation damage or low dose data collection, respectively.^1–4^ This technique not only determines the room temperature structure, but also enables the time-resolved molecular dynamics to be visualized through pump-probe experiments.^1–4^ Therefore, SX experiments allow for in-depth analyses of the more biologically reliable crystal structure of macromolecules beyond conventional X-ray crystallography using single crystal diffraction at cryogenic temperatures. In SX experiments, it is important to deliver the crystal sample serially and stably to the X-ray interaction point.^5^ Delivering the sample reliably not only reduces sample consumption but also enables data collection for accurate responses in time-resolved studies.^6^ Among the various sample delivery methods, including liquid jet injector,^7^ high viscosity injector,^8,9^ fixed-target scanning,^10–12^ and microfluidics,^13,14^ sample delivery using a viscous medium with an injector or syringe is widely applied to serial femtosecond crystallography (SFX) or serial millisecond crystallography (SMX), which can successfully determine structure using an XFEL or synchrotron, respectivley.^15–22^ The advantage of crystal delivery with a viscous delivery medium is the drastic reduction of sample consumption by lowering the flow rate of the sample loading from the injector or syringe.^22^

Based on their physical and chemical properties, sample delivery media can be categorized as either lipidic cubic phase (LCP),^15^ oil-based,^16–18^ or hydrogel-based.^17–21^ Among them, LCP and grease, which have hydrophobic characteristics, provide the most stable injection streams and are the most widely used delivery media in SX experiments.^15–18^ LCP, which is usually used with monoolein, is widely applied in the process of membrane protein crystallization, and its crystallized membrane protein embedded in LCP can be used as a sample delivery medium.^15,23^ The LCP delivery medium enables the sample to be delivered very steadily, even at flow rates of 0.001–0.3 μl/min.^9,24^ Moreover, LCP can be used as a sample delivery medium for soluble protein crystal samples, and it typically produces a stable injection stream at atmospheric pressure.^25^ The phase of LCP can be changed to the lamellar crystalline phase in a vacuum environment, but the addition of shorter chain lipids can prevent this phase transition.^15^ However, the phase of LCP can be transformed into lamellar, hexagonal, or sponge, depending on the crystallization or experimental environment, and it is not suitable for use with high concentration of ammonium sulfate.^9,26^ Oil-based grease matrices, such as mineral oil grease,^16^ synthetic grease Super Lube,^17^ and nuclear grade grease,^18^ also provide stable injection streams and have been applied to various SFX experiments. However, because this material is usually in a semisolid state at room temperature, it is difficult to transfer it to a syringe or other devices, which can cause sample loss during sample preparation. On the other hand, the stability of the injection stream for typical sample delivery media deteriorates depending on the physical or chemical reactions with the specific crystal sample or crystallization solution. Thus, it is very important to continuously develop various potential alternative delivery materials.^22^

Shortening consists of fats formulated from oils, and it is a very safe substance used primarily in food production.^27^ This material is readily available and has the advantage of being much less expensive than previously reported sample delivery media. In general, shortening has a solid state at room temperature and can be changed to a liquid state by raising its temperature, which is considered useful for sample preparation.

Here, I introduce the characterization and preparation of a shortening injection matrix for serial crystallography. The background scattering in shortening was negligible in data processing and showed similar or low scattering intensity compared to that of LCP. SMX experiments were performed using commercially available shortening, and the room temperature crystal structures of lysozyme and glucose isomerase were determined at resolutions of 1.5–2.0 Å. Shortening can be used for crystal delivery for SFX or SMX experiments.

## Results and Discussion

### Characterization and preparation of shortening injection matrix

In this experiment, two types of commercially available oil-based shortening were used, which are denoted as shortening A (composed of palm oil and tallow) and shortening B (composed of palm oils, tallow, and tocopherol). Commercially available shortening can exhibit physically varying melting points,^28^ which is due to the composition of the shortening, the proportions of oils, and the manufacturing process.^27^ To apply shortening to an injection matrix for an SX experiment, the melting temperatures of shortenings A and B were screened in the range of 20 to 40 °C. The results showed that the melting temperatures of shortenings A and B were approximately 28.5 and 26.5 °C, respectively.

Next, the sample preparation method using shortening was established for the SX experiment. The crystals embedded in shortenings A and B were prepared by mechanical mixing using a dual syringe setup (Figure 1). Because the melting temperature of the shortening is below typical body temperature, the liquid phase of the shortening can be achieved by holding the glass vial containing the shortening by hand, but this takes several minutes. To reduce sample preparation time, in this experiment, the glass vial containing the shortening was immersed in hot water (~100 °C) in a beaker for approximately 10–20 s to obtain its liquid phase (Figure 1a). The liquid phase shortening was transferred to a 100 μl syringe using the pipette and kept at room temperature or low temperature in an incubator until solidified (Figure 1a). The crystal suspension was transferred to another syringe using a pipette (Figure 1b). This syringe was then placed vertically and left for at least 10 minutes (see below). When the crystals sank to the bottom, the plunger was pushed upwards and the supernatant of the crystal suspension was removed, leaving the crystals (Figure 1b). Next, two syringes containing shortening and crystals were connected using a coupler and the sample was gently mixed by moving the plunger back and forth until the crystal sample was evenly distributed (Figure 1c). The mixture sample of crystals and shortening was transferred to a syringe, and the partner syringe with the coupler was removed. After connecting the syringe needle to syringe containing the mixture, the crystals embedded in the shortening were extruded from the syringe needle using a syringe pump (Figure 1d).

**Figure 1.**
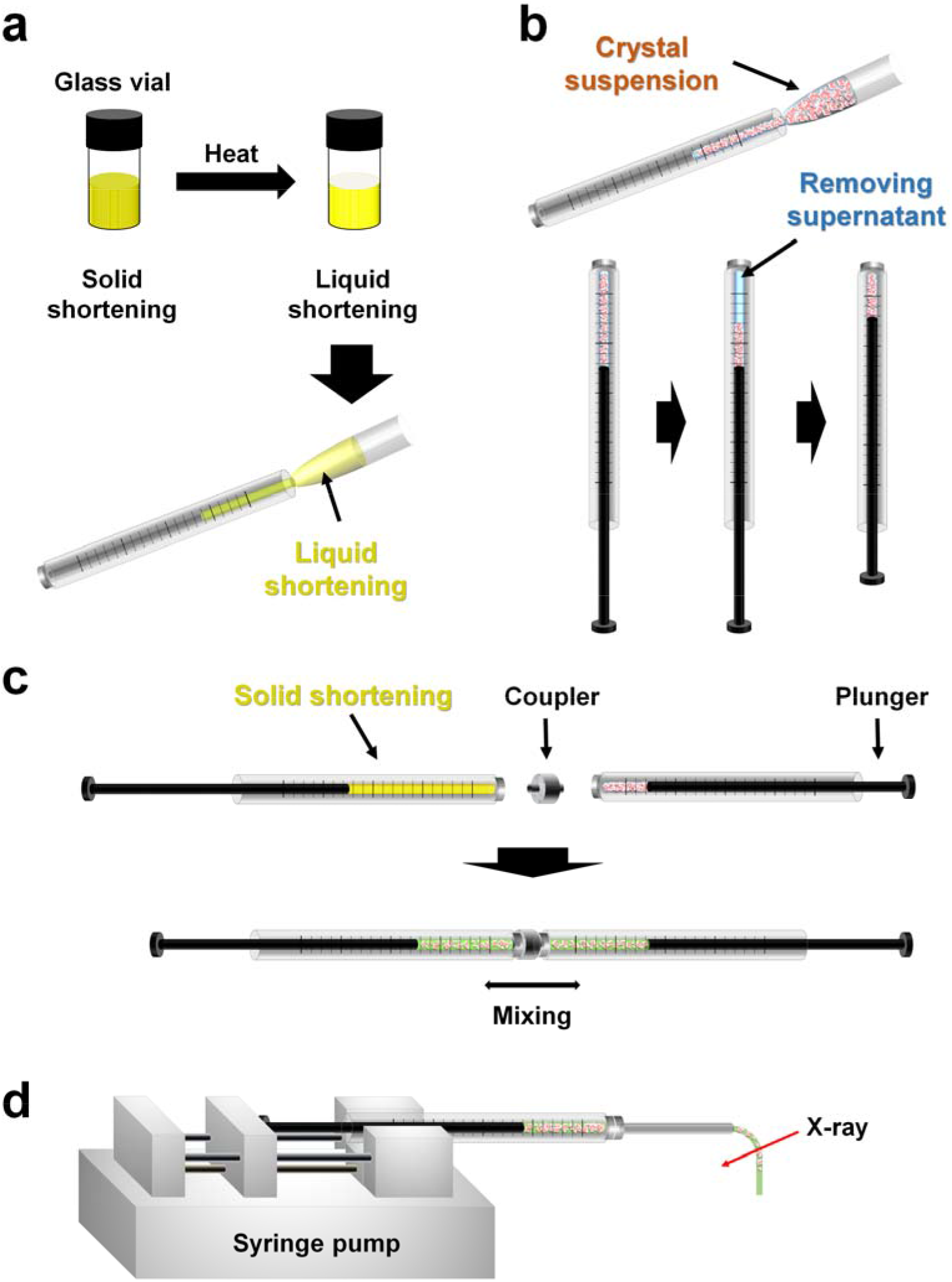
Preparation of shortening injection matrix. (a) Solid shortening in glass vial was immersed in hot water, and the shortening liquid was transferred to a syringe using a pipette.(b)Crystal suspension in syringe was stood vertically. When crystals settled, then crystal solution was removed. (c) The syringes containing the shortening and crystals were connected with a coupler and gently mixed. (d) The crystals embedded in the shortening were passed through the syringe needle using a syringe pump and delivered to the X-ray position.

### Screening the injection stream of shortening

To determine whether shortening can deliver the crystal sample for SX, stable injection streams for shortenings A and B were investigated using a commercially available syringe and syringe pump. The syringe containing the shortening was connected to the syringe needle with a 168 μm inner diameter (ID), and it was vertically installed into the syringe pump. The shortening was extruded from the syringe needle by pushing the syringe plunger using a syringe pump. At low flow rates of less than 200 nl/min (velocity: 9.02 mm/s), shortening provided a curled injection stream irregularly at the tip of the needle from which the sample came out. To solve this problem, the initial flow rate of the shortening was increased to 3 μl/min (135.33 mm/s) and the shortening was extruded for 3–5 s before stopping the syringe pump. The extruded shortening formed a stream of approximately 10 mm in length, which remained one stream due to its viscosity. When the shortening stream was directed downward in line with gravity, even when shortenings A and B were ejected at a flow rate lower than 200 nl/min (velocity: 4.51 mm/s), they flowed very steadily in the downward direction (Figures 2).

**Figure 2.**
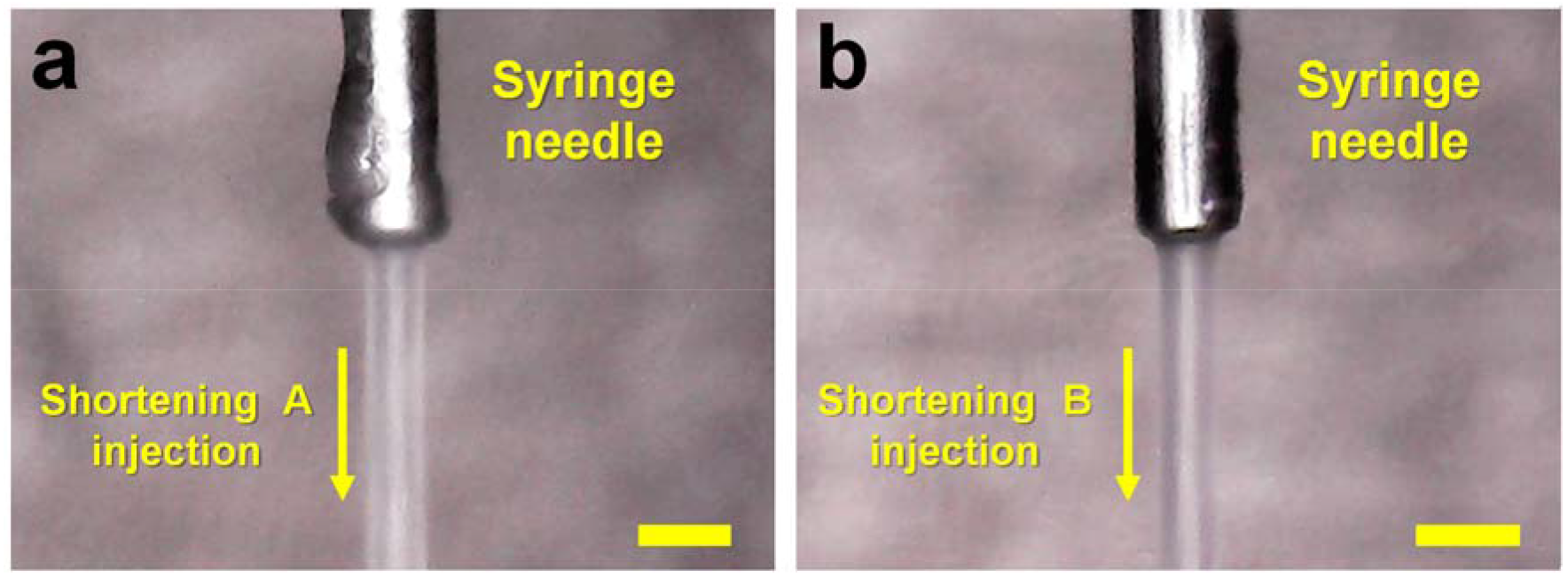
Snapshots of the injection streams of (a) shortenings A and (b) shortening B at a flow rate of 200 nl/min.

The main component of shortening is the hydrophobic lipid triglyceride, whereas the crystal solution is hydrophilic. Therefore, the shortening and crystal suspension have different polarities. To achieve a stable shortening injection stream, a suitable ratio of shortening to crystal suspension was investigated. In the mechanical mixing method using a dual syringe setup, the shortening was mixed with lysozyme crystal suspension at ratios of 5:5, 6:4, 7:3, 8:2, and 9:1. When the crystal solution was more than 30%, an unmixed crystal solution was ejected in the middle of the shortening injection stream and disrupted the continuous injection stream. By contrast, less than 20% of the crystal solution provided a stable shortening injection stream. However, chemically minimizing the crystal solution produced a more continuous injection stream. Therefore, a syringe containing a crystal sample was set up vertically before the dual mixing, and when the crystals had settled, the upper layer crystal solution was removed and then mixed with the shortening (Figure 1c). The percentage of settled crystal suspension was approximately 15–20% of the total, and the shortening containing crystals provided a steady and stable injection stream like LCP injection.

### Measurement of background scattering

The X-ray exposed sample delivery media generate background scattering, which can affect the signal-to-noise (SNR) ratio during data processing.^19,21^ The background scattering from the delivery medium is one of the most important criteria for delivery medium selection for SX experiments.^22^ The analyses of the background scattering of shortenings A and B were performed with the LCP delivery medium consisting of 60% (v/v) monoolein. The delivery media were extruded from the syringe needle with a 168 μm ID, and they were exposed to X-ray radiation with a photon flux of 1.3 × 10^12^ for 100 ms. For shortening A, five background scattering rings were observed at 3.90, 4.26, 4.41,14.32, and 44.02 Å (Figure 3a), giving average analog-digital units (ADU) values of 25, 37, 38, 21, and 155, respectively (Figure 3c). For shortening B, five background scattering rings were observed at 3.90, 4.26, 4.41,14.32, and 44.02 Å (Figure 3a), giving average ADU values of 11, 16, 15, 14, and 59, respectively (Figure 3c). Therefore, shortening A and shortening B show almost identical background scattering ring patterns in a similar resolution area, whereas the background intensity of shortening B was 2-3 times lower than that of shortening A (Figure 3c). The scattering ring patterns from shortenings A and B are considered to be the scattering from the lipid packing in the solid phase. For LCP, two background scattering rings were observed at 4.5 Å and 25–100 Å, and the average ADU values were 20 and 110, respectively (Figure 3c). Analysis of the 2D profile of background scattering intensity revealed that shortening A had slightly higher background scattering intensity than LCP at approximately 45 Å and 3.0–5.5 Å (Figure 3c). On the other hand, shortening B showed lower overall intensities of background scattering than LCP, excluding the intensity of the scattering ring at approximately 14 Å (Figure 3c). Therefore shortening can have different background scattering depending on its production process and composition, and the shortening used in this study showed similar or slightly lower background scattering compared to LCP.

**Figure 3.**
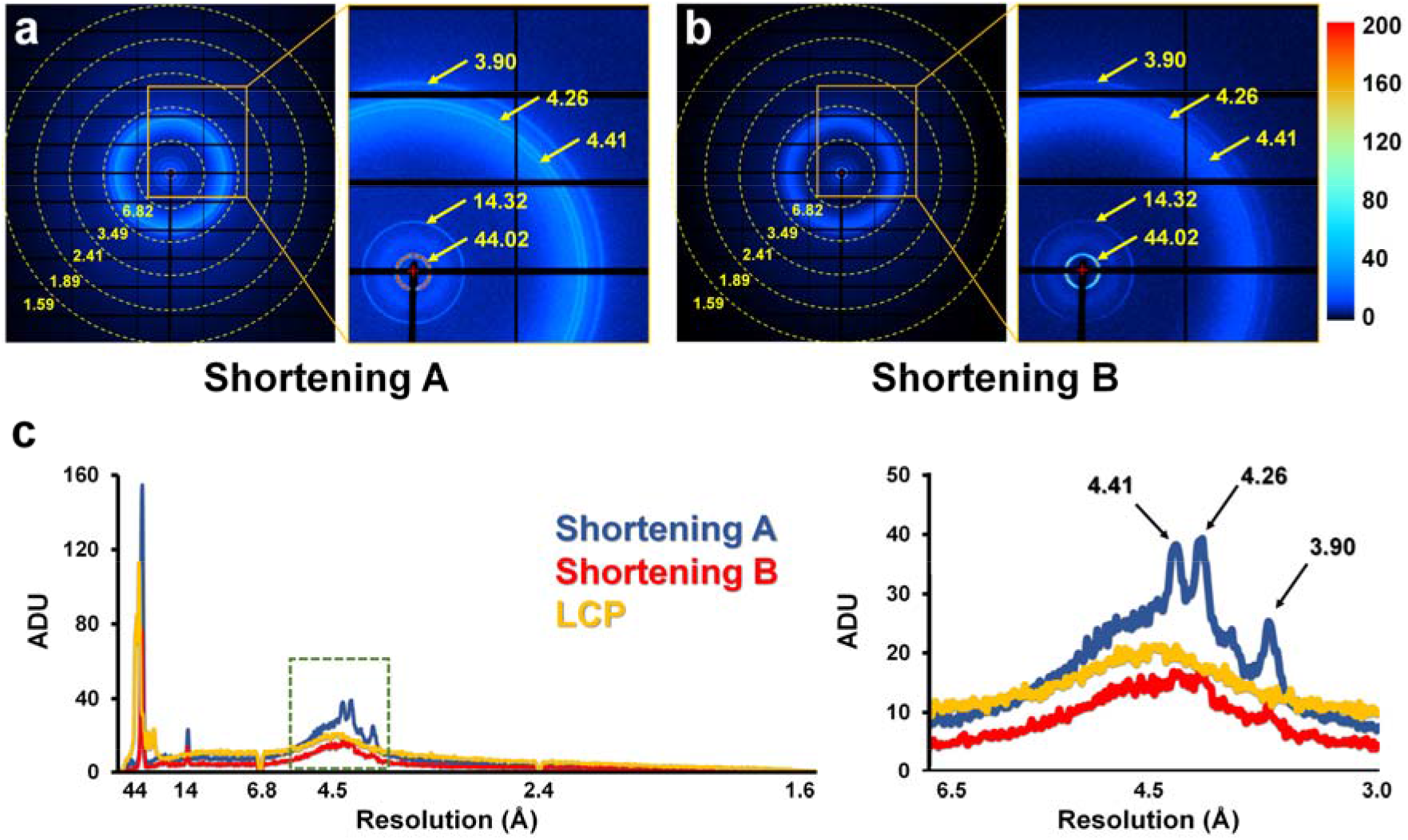
Background X-ray scattering of shortening: (a) shortening A and (b) shortening B.2D profile of the scattering intensities of shortening A (blue), shortening B (red), and LCP (yellow). (right) close-up view of green dotted box.

### SMX using the shortening injection matrix

To demonstrate the application of a shortening injection matrix, SMX experiments using shortenings A and B were performed with the crystals of lysozyme and glucose isomerase as model samples. During sample preparation, the crystallization solution from crystal suspension was removed as much as possible, and mechanical mixing was performed with shortenings A or B using dual-syringe setups. The proportion occupied by crystals was less than 20% of the total sample volume. In these experiments, the syringe containing the crystals embedded in shortening was installed on the syringe pump in the horizontal direction to avoid beamline interference. The crystal samples were extruded from the syringe needle with a 168 μm ID at a flow rate of 200–300 nl/min using the syringe pump. In an offline injection study, the shortening injection stream was stably extruded from the syringe at room temperature, but the extruded injection stream of shortening in the experimental hutch at 25 °C was less viscous like the unsolidified form, making it the injection stream unstable. It is believed that the shortening was partially melted by heat from the beamline device or the camera light. To solve this problem, the internal temperature of the experimental hutch was kept at 20 °C, which provided a very stable shortening injection stream. For shortening A, totals of 64000 and 48000 images were collected for glucose isomerase and lysozyme, respectively. There were 13651 and 15643 final indexed images of glucose isomerase and lysozyme, respectively (Supplementary Figures S1 and S2). Glucose isomerase delivered in shortening A was processed up to 1.9 Å, and the overall SNR, CC, and R_split_ were 4.09, 0.9576, and 19.11, respectively. Lysozyme delivered in shortening A was processed up to 1.8 Å, and the overall SNR, CC, and R_split_ were 100, 7.87, 0.9937, and 7357, respectively. The final R_work_/R_free_ values of glucose isomerase and lysozyme in shortening A were 17.95/21.43 and 17.66/20.53, respectively. For shortening B, totals of 48000 images were collected for glucose isomerase and lysozyme, respectively. There were 16522 and 27413 final indexed images of glucose isomerase and lysozyme in shortening B, respectively (Supplementary Figures S3 and S4). Glucose isomerase delivered in shortening B was processed up to 1.9 Å, and the overall SNR, CC, and R_split_ were 4.10, 0.9669, and 17.88, respectively. Lysozyme delivered in shortening B was processed up to 1.5 Å, and the overall SNR, CC, and R_split_ were 9.27, 0.9936, and 6.58, respectively. The R_work_/R_free_ values of glucose isomerase and lysozyme derived in shortening B were 17.56/20.10 and 18.49/20.69, respectively.

The glucose isomerase delivered in shortenings A and B showed clear electron density maps for amino acids from Tyr3 to Arg387 (Figure 4a). The active site of glucose isomerase contains two metal binding sites for substrate binding and catalytic function.^29,30^ These metal binding sites of glucose isomerase are well-defined in the electron density map, and there was no negative fo-fc electron density map counted at 3 σ, indicating there was no significant radiation damage (Supplementary Figure S5). The glucose isomerase structures delivered in shortenings A and B had high similarity with previously reported glucose isomerase structures delivered using grease (PDB code 4W4Q)^31^ and nylon mesh-based fixed target scanning (6IRK)^12^ with a root mean squared deviation of 0.1594–0.2549 for Cα atoms (Supplementary Figure S6).

**Figure 4.**
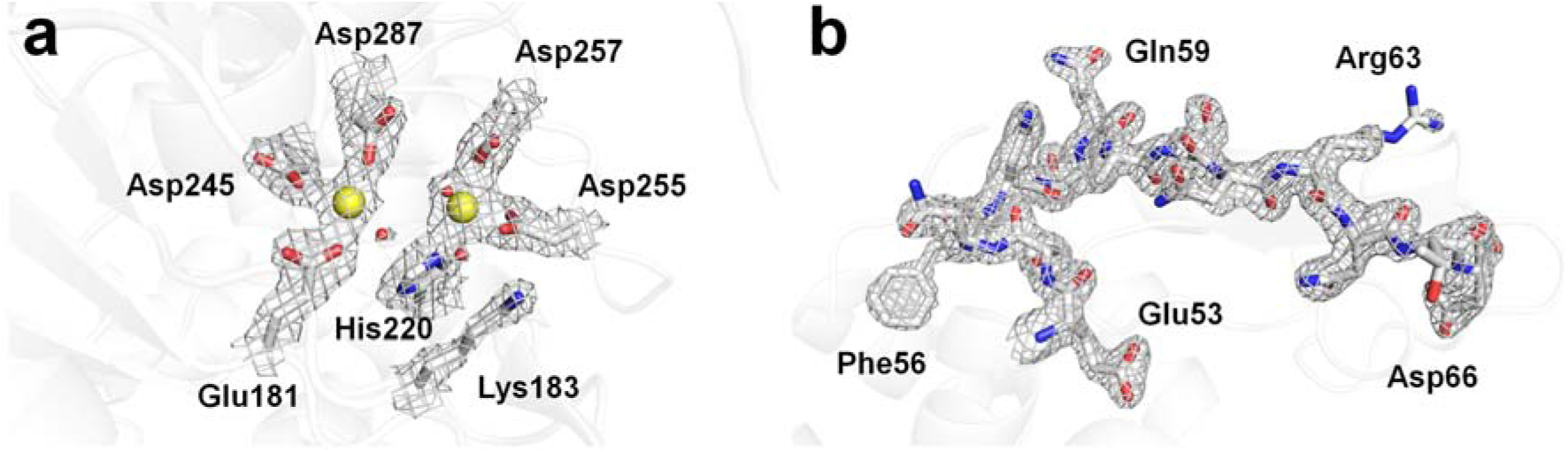
2Fo-Fc electron density maps of (a) glucose isomerase (light blue mesh, 1.2 σ) delivered in shortening A and (b) lysozyme (light blue mesh, 1.5 σ) delivered in shortening B.

**Table 1.** Data collection and refinement statistics.

| Data collection | Shortening A |  | Shortening B |  |
| --- | --- | --- | --- | --- |
|  | GI | Lysozyme | GI | Lysozyme |
| Wavelength (Å) |  | 0.9796 |  |  |
| Photons/sec<sup>a</sup> |  | ~ 1.3 x 10<sup>12</sup> |  |  |
| Expose time |  | 100 ms |  |  |
| Space group | I222 | P4<sub>3</sub>2<sub>1</sub>2 | I222 | P4<sub>3</sub>2<sub>1</sub>2 |
| Cell dimensions (Å) |  |  |  |  |
| *a* | 94.19 | 79.55 | 94.19 | 79.18 |
| *b* | 99.92 | 79.55 | 99.92 | 79.18 |
| *c* | 103.25 | 38.51 | 103.25 | 38.34 |
| No. collected images | 64000 | 48000 | 48000 | 48000 |
| No. of hits | 25626 | 18926 | 30418 | 29290 |
| No. of indexed images | 13651 | 15643 | 16522 | 27413 |
| Resolution (Å) | 72.46-1.90 | 80.00-1.80 | 72.46-2.00 | 80.00-1.50 |
|  | (1.96-1.90) | (1.86-1.80) | (2.07-2.00) | (1.55-1.50) |
| Unique reflections | 38725 (3832) | 11974 (1148) | 33248 (3247) | 20406 (1995) |
| Completeness | 100.0 (100.0) | 100.0 (100.0) | 100.0 (100.0) | 100.0 (100.0) |
| Redundancy | 426.9 (292.6) | 1451.9 (1011.7) | 701.6 (481.7) | 756.6 (285.8) |
| *I*/$\sigma$(*I*) | 4.09 (2.07) | 7.87 (2.21) | 4.10 (1.72) | 9.27 (1.68) |
| *R*<sub>split</sub><sup>b</sup> | 19.11 (53.97) | 7.57 (46.32) | 17.88 (63.03) | 6.58 (70.95) |
| CC | 0.9576 (0.6392) | 0.9937 (0.8016) | 0.9669 (0.5778) | 0.9936 (0.5183) |
| CC* | 0.9891 (0.8831) | 0.9984 (0.9433) | 0.9915 (0.8558) | 0.9984 (0.8262) |
| Wilson B factor (Å<sup>2</sup>) | 30.85 | 36.24 | 31.57 | 39.72 |
| **Refinement statistics** |  |  |  |  |
| Resolution (Å) | 71.80-1.90 | 56.18-1.80 | 71.80-2.0 | 55.98-1.50 |
| *R*<sub>factor</sub> / *R*<sub>free</sub> (%)<sup>c</sup> | 17.95/21.43 | 17.66/20.53 | 17.56/20.10 | 18.49/20.69 |
| B-factor (Averaged) |  |  |  |  |
| Protein | 34.02 | 36.88 | 33.63 | 40.33 |
| Ligands | 33.50 | 36.45 | 22.90 | 44.35 |
| R.m.s. deviations |  |  |  |  |
| Bond lengths (Å) | 0.006 | 0.006 | 0.007 | 0.006 |
| Bond angles (°) | 0.825 | 0.787 | 0.831 | 0.797 |
| Ramachandran plot (%) |  |  |  |  |
| favored | 96.86 | 98.43 | 97.12 | 99.21 |
| allowed | 2.88 | 1.57 | 2.62 | 0.79 |
| outliers | 0.26 |  | 0.26 |  |

Highest resolution shell is shown in parentheses.
<sup>a</sup> Sample position
<sup>c</sup> $R_{work} = \Sigma ||F_{obs}| - |F_{calc}|| / \Sigma |F_{obs}|$ , where $F_{obs}$ and $F_{calc}$ are the observed and calculated structure-factor amplitudes respectively. $R_{free}$ was calculated as $R_{work}$ using a randomly selected subset (10%) of unique reflections not used for structure refinement.

The lysozyme delivered in shortenings A and B showed clear electron density maps for amino acids from Lys19 to Leu147 (Figure 4b). In lysozyme delivered in shortening A, four disulfide bonds (Csy24-Csy145, Csy48-Csy133, Cys82-Cys98, and Csy94-Csy112) were well-defined, and there was no negative fo-fc electron density map counted at 3 σ, indicating that there was no significant radiation damage (Supplementary Figure S7a). In contrast, in shortening B, lysozyme delivered three disulfide bonds (Csy24-Csy145, Csy48-Csy133, and Cys82-Cys98) with a partial negative fo-fc electron density map counted at 3 σ (Supplementary Figure S7b). Because the procedure and experimental setup of the SMX experiments using shortening A and B were almost identical, the partial negative fo-fc map of disulfide of lysozyme delivered in shortening B is presumed to be influenced by the delivery material rather than radiation damage caused by the X-rays. On the other hand, the overall lysozyme structures determined using both shortening A and B are similar to previously reported lysozyme structures as delivered by liquid jet sample injector (PDB code 4ET8)^31^, droplet injector (5DM9)^32^ and polyacrylamide (6IG6) with a root mean squared deviation of 0.0640–0.1496 for whole Cα atoms (Supplementary Figure S8).

## Conclusion

Here, I reported the characterization and preparation of shortening as a sample delivery medium for SX experiments. SMX was successfully demonstrated using two shortening delivery media. However, shortening may require a screening procedure before in can be used, as it may exhibit differences in melting temperature, background scattering, and reduction of disulfide bonds depending on its production process and composition. In addition, because shortening is mostly fat, lipid-related protein samples may be unstable due to specific or non-specific interactions between crystals and shortening. Nevertheless, the advantages of shortening are clear in that it is easy to store and much less expensive than previously reported sample delivery media (e.g., the price is 45,000,000 times lower than that of LCP). Moreover, the transferal of shortening is straightforward by the temperature control of the solid-liquid phase. Therefore, shortening is a promising delivery material for SX experiments with existing hydrophobic delivery materials.

## Materials and Methods

### Crystallization

Glucose isomerase from *Streptomyces rubiginosus* and lysozyme from chicken egg whites were purchased from Hampton Research (HR7-098) and Sigma-Aldrich (L6876), respectively. Glucose isomerase was supplied as a crystal suspension, and it was used directly for SMX experiments, as previously reported.^12^ The lysozyme was crystallized using a previously reported procedure.^12^ The crystal sizes of glucose isomerase and lysozyme were 30–40 μm and <60 μm, respectively.

### Characterization of shortening

Shortening A was purchased from Samyang (Republic of Korea) and shortening B from Ottogi (Republic of Korea). Shortening A was composed of palm oil and beef tallow. Shortening B was composed of palm olein oil, palm stearin oil, palm hydrogenated oil, tallow, and d-Tocopherol. The melting temperatures of shortenings A and B were measured on a water bath and judged visually. The solid phase shortening at room temperature was initially transferred to a glass vial using a spatula. During the transfer of the shortening to the syringe, the shortening in the glass vial was immersed in hot water (~100 °C) in a beaker for 10**–**20 s. Liquid shortening was transferred into a 100 μl syringe (Hamilton, 81065-1710RNR) using a pipette, and was then left to stand until solid. A syringe needle of 168 ID was connected to the syringe containing the shortening, and the syringe was vertically installed in a Fusion Touch 100 syringe pump (CHEMYX). The syringe plunger was pushed by a mechanical force from the syringe pump^33^ and extruded the sample at a flow rate of <200 nl/min.

### Crystal embedding in shortening

Solid shortening in glass vials was dissolved by soaking in hot water (> 100 °C) for 20 seconds. The shortening solution (50 μl) was transferred to a 100 μl syringe and stored at room temperature until it reached a solid state. The crystal suspension (20 μl) was transferred to a 100 μl syringe. This syringe was vertically orientated for 10 min. When crystals settled on the bottom, the supernatant was removed using a pipette. The syringes containing the shortening and crystals were connected using a syringe coupler and mixed with the plunger gently moving back and forth more than 30 times. The mixture sample was transferred to a syringe and the emptied partner syringe with the coupler was removed. The syringe containing the crystals embedded shortening was connected with a syringe needle of 168 μm ID for SMX experiments.

### Data collection

SMX experiments using a shortening injection matrix were performed at the 11C beamline at Pohang Accelerator Laboratory (Republic of Korea). The temperature and humidity were 20 °C and 20%, respectively. The X-ray beam size focused by a Kirkpatrick-Baez mirror was approximately 4 (vertical) × 8 (horizontal) μm^2^ (FWHM) at the sample position. The photon flux at the sample position was 1.3 × 10^12^ photons/s, and the X-ray energy was 12.657 keV. The shortening containing crystals was delivered by a syringe pump-based sample delivery system^33^ through a syringe needle of 168 μm ID at a flow rate of 200-300 nl/min. Crystals were X-ray exposed for 100 ms. Diffraction images were recorded on a Pilatus 6M with 10 Hz readout.

### Data processing and structure determination

The hit images with diffraction pattern were filtered using the Cheetah program with the peakfinder8 algorithm.^34^ The hit images were indexed and merged using the CrystFEL program.^35^ The phase problem of the lysozyme and glucose isomerase was solved by molecule replacement using the Phaser-MR in PHENIX^36^, with the crystal structures of lysozyme (PDB code 6IG6)^21^ and glucose isomerase (PDB code 5ZYD)^30^ used as the search models. The model building and refinement were conducted using Coot^37^ and Phenix.refinement in PHENIX, respectively.^36^ The geometry was analysed using MolProbity.^38^ Figures were generated using PyMOL (https://pymol.org/).

### Background scattering analysis

Shortening A, shortening B, and LCP delivery media were passed through a syringe needle with a 168 μm ID. Each delivery medium was exposed to X-rays of 1.3 × 10^12^ photons/flux for 100 ms. The background scattering from 1.6 Å to the beam centre was analysed using the average intensity for 100 images. The background intensities of the delivery media were analysed using ADXV (https://www.scripps.edu/tainer/arvai/adxv.html).

## Ackowledgments

I thank the beamline staff at 11C beamline at Pohang Accelerator Laboratory for their assistance with data collection. The authors thank Global Science experimental Data hub Center (GSDC) at Korea Institute of Science and Technology Information (KISTI) for computational support. This work supported by a Korea University Grant. This work was funded by the National Research Foundation of Korea (NRF-2017M3A9F6029736).

## Author Contributions

K.H.N. performed all experiment and wrote the manuscript.

## Accession codes

The coordinates and structure factors have been deposited in the Protein Data Bank under the accession code 6KCA (Glucose isomerase-shortening A), 6KCB (Lysozyme-shortening A), 6KCC (Glucose isomerase-shortening B), 6KCD (Lysozyme-shortening B).

## Supplementary Data

**Supplementary Figure S1.**
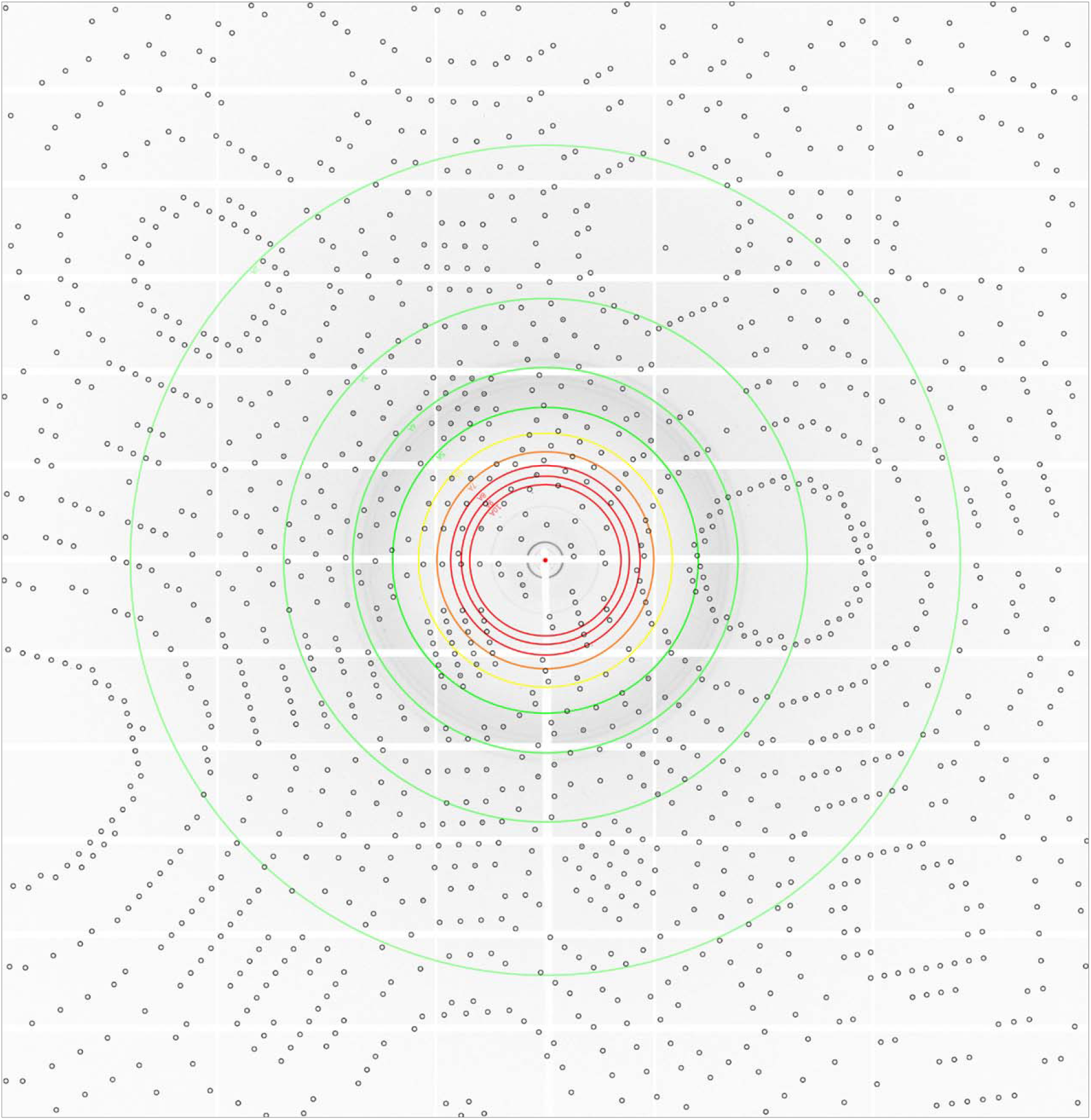
Indexed imaged of glucose isomerase delivered in shortening A.

**Supplementary Figure S2.**
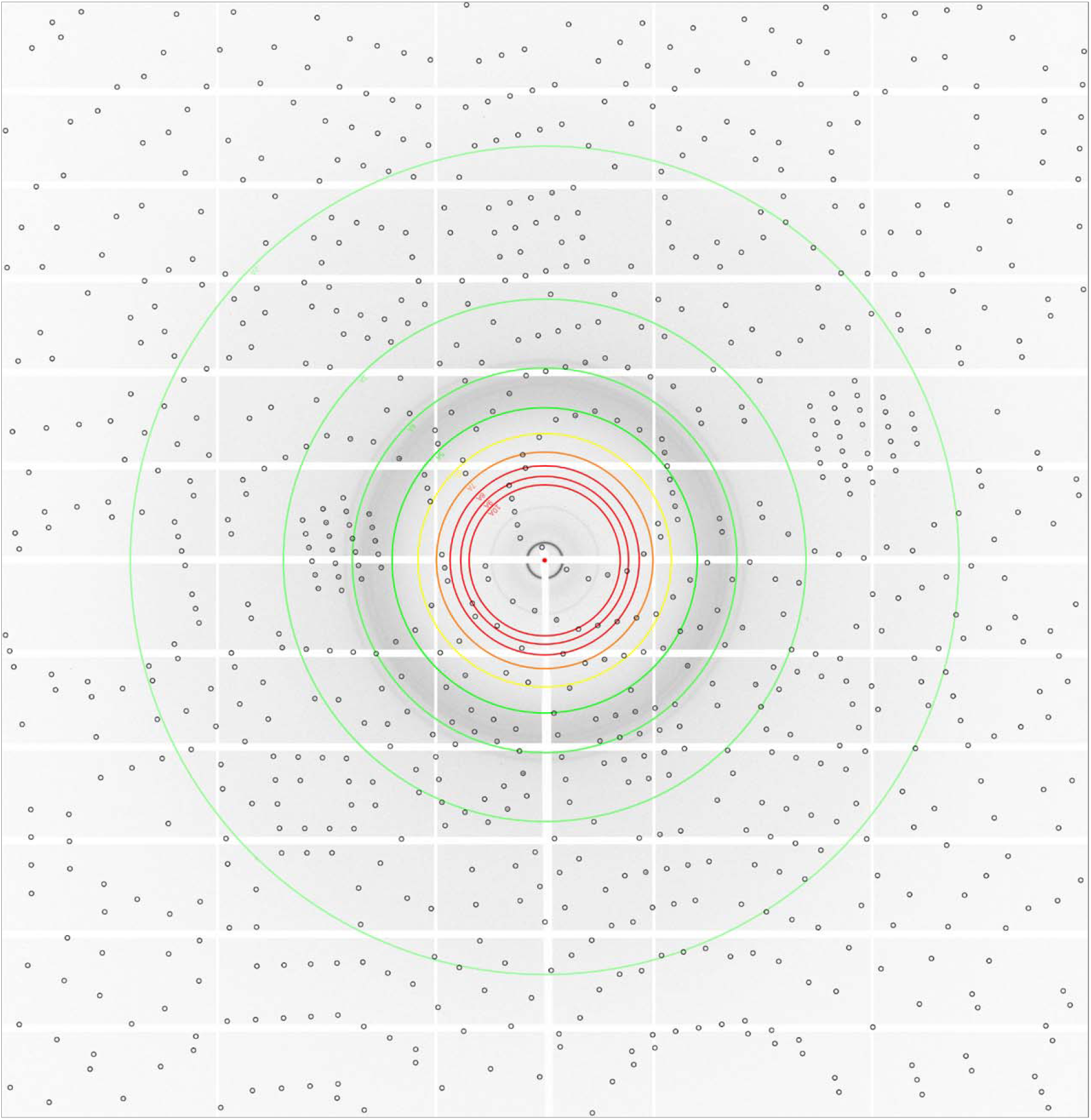
Indexed imaged of lysozyme delivered in shortening A.

**Supplementary Figure S3.**
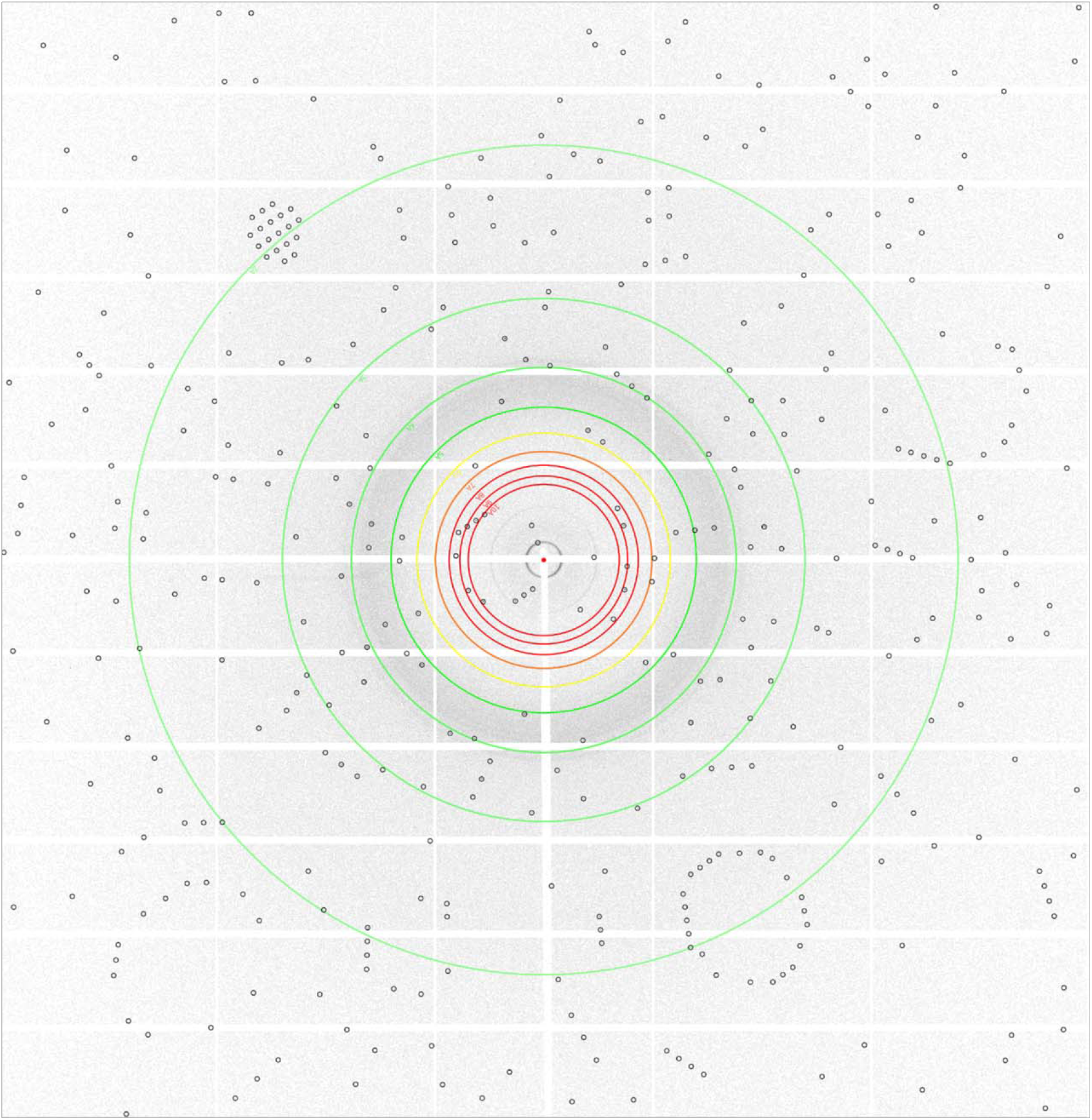
Indexed imaged of glucose isomerase delivered in shortening B.

**Supplementary Figure S4.**
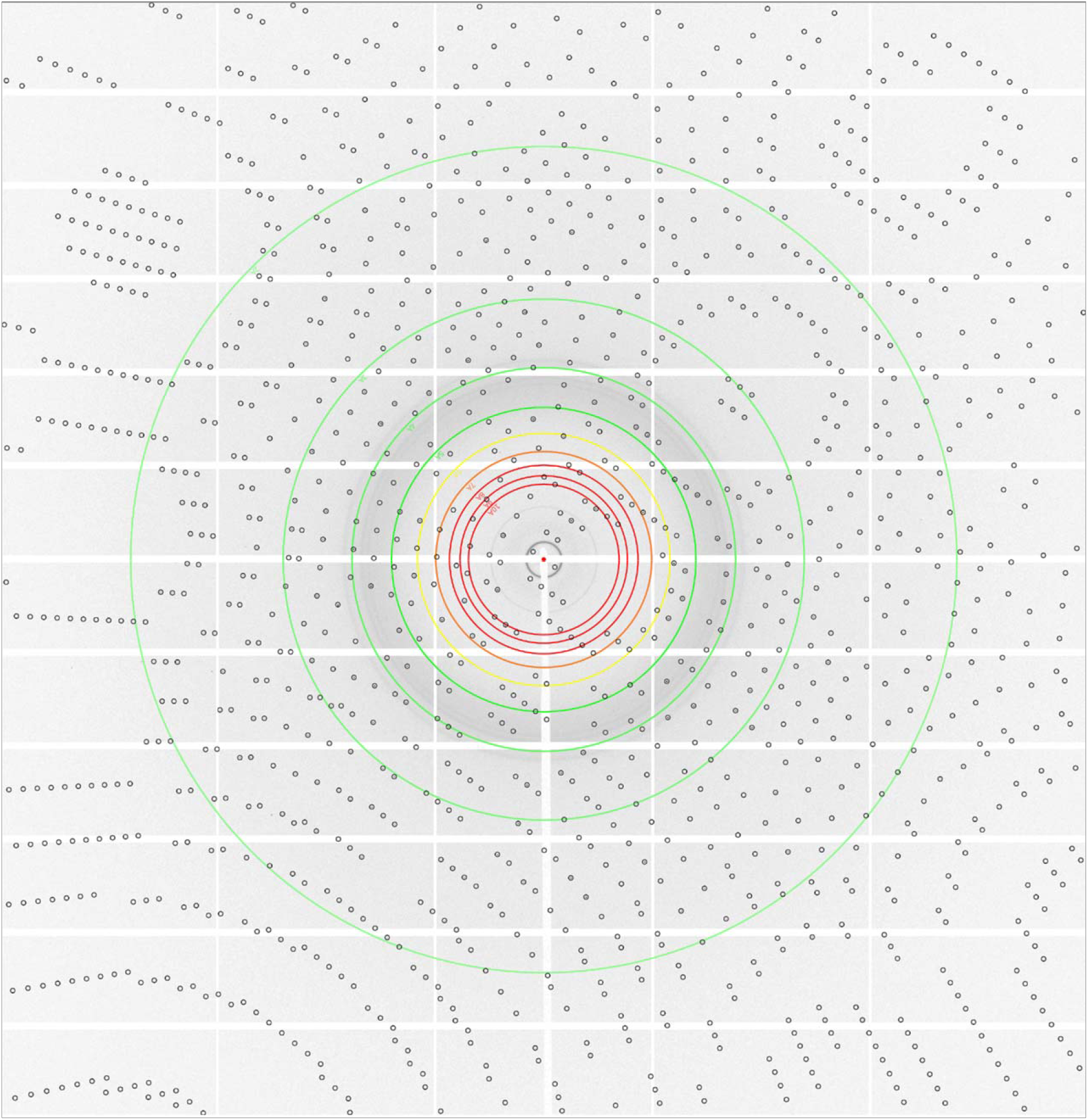
Indexed imaged of lysozyme delivered in shortening B.

**Supplementary Figure S5.**
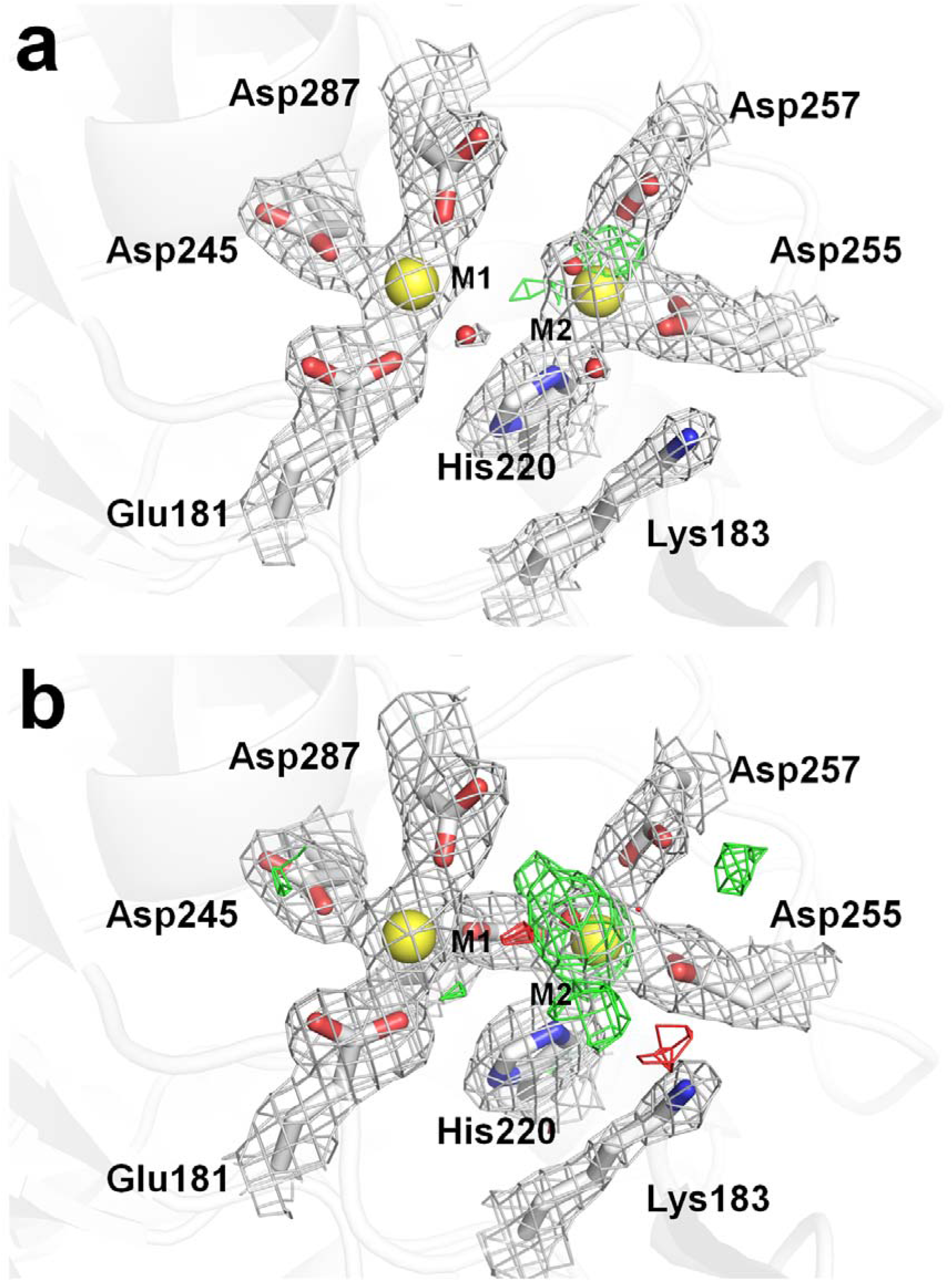
2Fo-Fc electron density map (grey, counted 1.5 σ) and Fo-Fc electron density map (green, counted 3 σ; red, counted - 3 σ) of active site of glucose isomerase delivered in (a) shortening A and (b) shortening B.

**Supplementary Figure S6.**
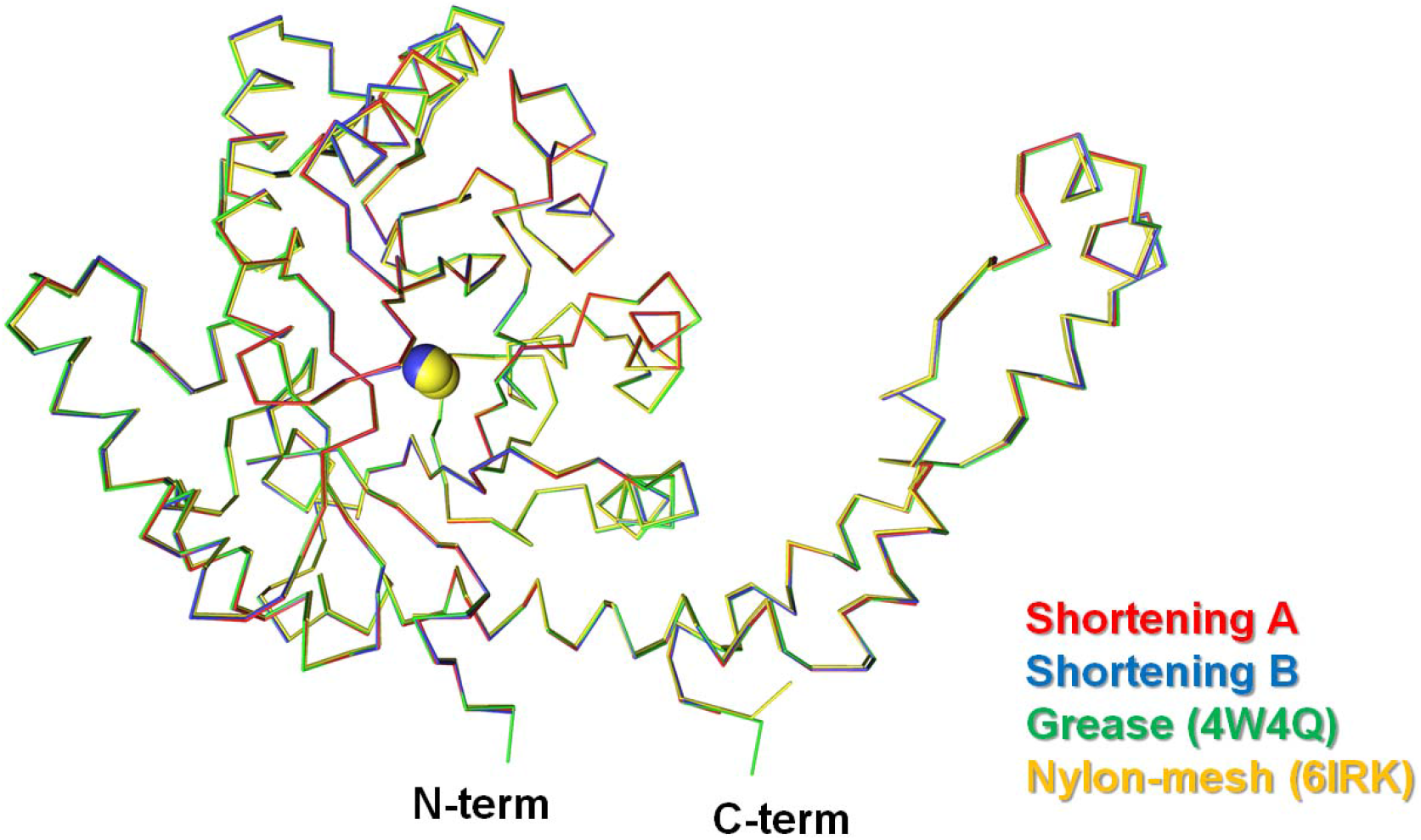
Superimposition of the crystal structure of glucose isomerase delivered in shortening A (red) and B (blue) with glucose isomerase delivered as grease delivery medium (PDB code: 4W4Q, green) and nylon-mesh based fixed-target scanning (6IRK, yellow).

**Supplementary Figure S7.**
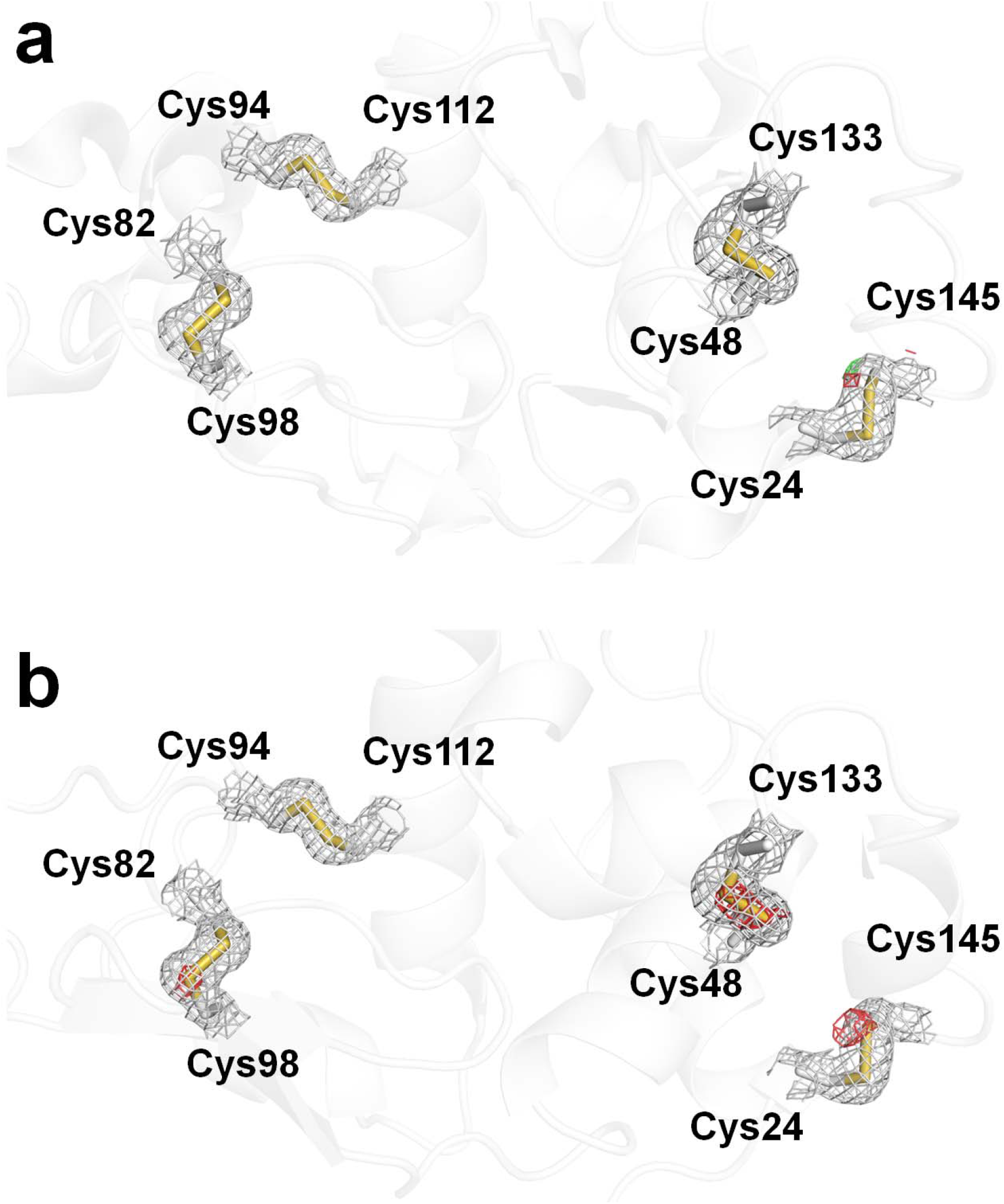
2Fo-Fc electron density map (grey, counted 1.5 σ) and Fo-Fc electron density map (green, counted 3 σ; red, counted - 3 σ) of disulfide bonds of lysozyme delivered in (a) shortening A and (b) shortening B.

**Supplementary Figure S8.**
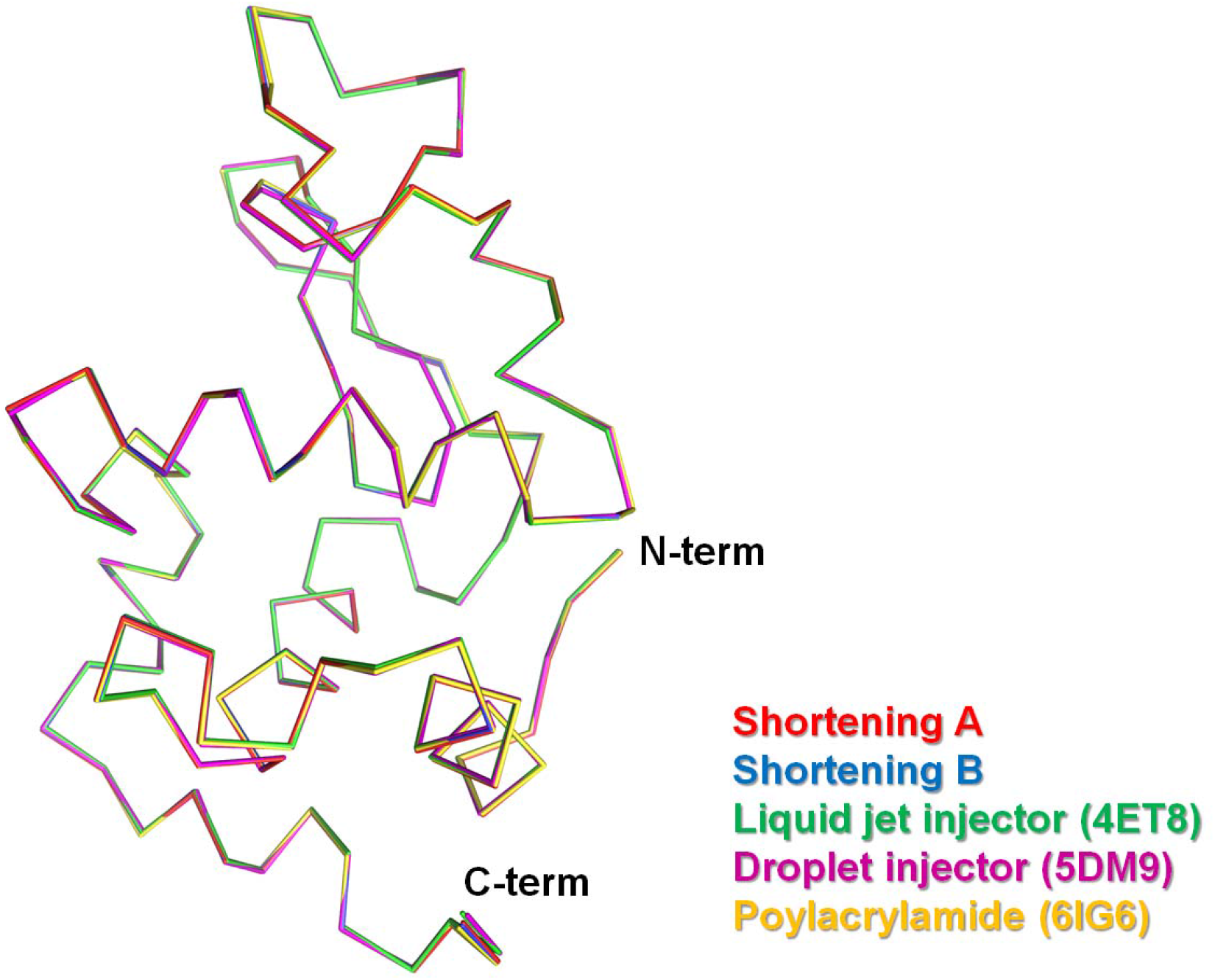
Superimposition of the crystal structure of lysozyme delivered in shortening A (red) and B (blue) with lysozyme delivered as liquid jet with Gas Dynamic Virtual Nozzle (PDB code: 4ET8, green), droplet injector (5DM9, purple) and polyacrylamide (6IG6, yellow).

